# Evaluating reproducibility of AI algorithms in digital pathology with DAPPER

**DOI:** 10.1101/340646

**Authors:** Andrea Bizzego, Nicole Bussola, Marco Chierici, Marco Cristoforetti, Margherita Francescatto, Valerio Maggio, Giuseppe Jurman, Cesare Furlanello

## Abstract

Artificial Intelligence is exponentially increasing its impact on healthcare. As deep learning is mastering computer vision tasks, its application to digital pathology is natural, with the promise of aiding in routine reporting and standardizing results across trials. Deep learning features inferred from digital pathology scans can improve validity and robustness of current clinico-pathological features, up to identifying novel histological patterns, e.g. from tumor infiltrating lymphocytes. In this study, we examine the issue of evaluating accuracy of predictive models from deep learning features in digital pathology, as an hallmark of reproducibility. We introduce the DAPPER framework for validation based on a rigorous Data Analysis Plan derived from the FDA’s MAQC project, designed to analyse causes of variability in predictive biomarkers. We apply the framework on models that identify tissue of origin on 787 Whole Slide Images from the Genotype-Tissue Expression (GTEx) project. We test 3 different deep learning architectures (VGG, ResNet, Inception) as feature extractors and three classifiers (a fully connected multilayer, Support Vector Machine and Random Forests) and work with 4 datasets (5, 10, 20 or 30 classes), for a total 53000 tiles at 512 × 512 resolution. We analyze accuracy and feature stability of the machine learning classifiers, also demonstrating the need for random features and random labels diagnostic tests to identify selection bias and risks for reproducibility. Further, we use the deep features from the VGG model from GTEx on the KIMIA24 dataset for identification of slide of origin (24 classes) to train a classifier on 1060 annotated tiles and validated on 265 unseen ones. The DAPPER software, including its deep learning backbone pipeline and the HINT (Histological Imaging - Newsy Tiles) benchmark dataset derived from GTEx, is released as a basis for standardization and validation initiatives in AI for Digital Pathology.

**Author summary:** In this study, we examine the issue of evaluating accuracy of predictive models from deep learning features in digital pathology, as an hallmark of reproducibility. It is indeed a top priority that reproducibility-by-design gets adopted as standard practice in building and validating AI methods in the healthcare domain. Here we introduce DAPPER, a first framework to evaluate deep features and classifiers in digital pathology, based on a rigorous data analysis plan originally developed in the FDA’s MAQC initiative for predictive biomarkers from massive omics data. We apply DAPPER on models trained to identify tissue of origin from the HINT benchmark dataset of 53000 tiles from 787 Whole Slide Images in the Genotype-Tissue Expression (GTEx) project. We analyze accuracy and feature stability of different deep learning architectures (VGG, ResNet and Inception) as feature extractors and classifiers (a fully connected multilayer, SVMs and Random Forests) on up to 20 classes. Further, we use the deep features from the VGG model (trained on HINT) on the 1300 annotated tiles of the KIMIA24 dataset for identification of slide of origin (24 classes). The DAPPER software is available together with the HINT benchmark dataset.

## Introduction

The rise of artificial intelligence (AI) methods for health data has soon crossed paths with disease complexity: patient cohorts are most frequently an heterogeneous group of subtypes diverse for disease trajectories, with highly variable phenotypes characteristics in terms of phenotypes (e.g. bioimages by radiology or pathology), response to therapy, clinical course, thus a challenge for machine-learning based prognoses. Nevertheless, the increased availability of massive annotated medical data from health systems and a rapid progress of machine learning frameworks has led to high expectations about the impact of AI on challenging biomedical problems [1].

In particular, Deep Learning has overcome *ad-hoc* pattern recognition methods in the analysis of Medical Images (MI), getting close to expert accuracy in the diagnosis of skin lesions [2, 3] or classification of colon polyps [4, 5], and expanding diagnostic options in radiomics [6] and in many other areas. Deep learning refers to a class of machine learning methods that model high-level abstractions in data through the use of modular architectures, typically composed by multiple nonlinear transformations estimated by training procedures. Notably, deep learning architectures based on Convolutional Neural Networks (CNNs) hold state-of-the-art accuracy in numerous image classification tasks without prior feature selection. Further, intermediate steps in the pipeline of transformations implemented by CNNs or other deep learning architectures can provide a mapping (*embedding*) from the original feature space into a *deep feature* space. Of interest for medical diagnosis, deep features can be used for interpretation of the model and can be directly employed as inputs to other machine learning models.

Deep learning methods have been applied to analysis of histological images for diagnosis and prognosis. Mobadersany and colleagues [7] combine in the SCNN architecture a deep learning CNN with traditional survival models to learn survival-related patterns from histology images, predicting overall survival of patients diagnosed with gliomas. Predictive accuracy of SCNN is comparable with manual histologic grading by neuropathologists. Further, by incorporation of genomic variables for gliomas in the model, the extended model significantly outperforms the WHO paradigm based on genomic subtype and histologic grading. Similarly, deep learning models have been successfully applied to histology for colorectal cancer [8], gastric cancer [9] and breast cancer [10].

As human assessments of histology are subjective and hard to repeat, computational analysis of histology imaging within the information environment generated from a digital slide (digital pathology) and advances in scanning microscopes have already allowed pathologists to gain a much more effective diagnosis capability and dramatically reduce time for data sharing. Starting from the principle that underlying differences in the molecular expressions of the disease may manifest as tissue architecture and nuclear morphologic alterations [11], it is clear that automatic detection of disease aggressiveness and patient subtyping has a key role aiding therapy in cancer and other diseases. Digital pathology is in particular a key tool for the immunotherapy approach, which stands on the characterization of Tumor Infiltrating Lymphocytes (TILs) [12].

Indeed, quantitative analysis of the immune microenvironment by histology is crucial for personalized treatment of cancer [13, 14], with high clinical utility of TILs assessment for risk prediction models, adjuvant, and neoadjuvant chemotherapy decisions, and for developing the potential of immunotherapy [15, 16]. Digital pathology is thus a natural application for machine learning, with the promise of aiding in routine reporting and standardizing results across trials. Notably, deep learning features learned from digital pathology scans can improve validity and robustness of current clinico-pathological features, up to identifying novel histological patterns, e.g. from TILs.

On the technical side, usually deep learning models for digital pathology are built upon architectures originally aimed at tasks in other domains and trained on non-medical datasets. This is a foundational approach in machine learning, known as *transfer learning.* Given domain data and a network pre-trained to classify on huge generic databases (e.g. ImageNet, with over 14 million items and 20 thousand categories), there are three basic options for transfer learning, i.e. to adapt the classifier to the new domain: a) train a new machine learning model on the features preprocessed by the pre-trained network from the domain data; b) retrain only the deeper final layers (the *domain layers*) of the pre-trained network; c) re-train the whole network starting from the pre-trained state. A consensus about the best strategy to use for medical images is still missing [17, 18].

In this study we aim to address the issue of reproducibility and validation of machine learning models for digital pathology. Reproducibility is a burning concern in biomarker research [19], and in science in general [20], with scientific communities, institutions, industry, and publishers struggling to foster adoption of best practices, with initiatives ranging from enhancing reproducibility of high-throughput technologies [21] to improving the overall reuse of scholarly data and analytics solutions (*e.g.* the FAIR Data Principles [22]). As an example, the MAQC initiative [23, 24], is led by the US FDA to investigate best practices and causes of variability in the development of biomarkers and predictive classifiers from massive omics data (e.g. microarrays, RNA-Seq or DNA-seq data) for precision medicine. The MAQC projects adopt a Data Analysis Plan (DAP) that force bioinformatics teams to submit classification models, top features ranked for importance and estimate of performance all built on training data only, before testing on unseen validation data. This approach helps mitigating the risk of selection bias in complex learning pipelines, where the bias can be born in one of many preprocessing steps as well as in the downstream machine learning model. Further, it clarifies that increasing task difficulty is often linked to a decrease in accuracy measures and also of stability of the biomarker lists [25].

Although openness in sharing algorithms and benchmark data is a solid attitude of the machine learning community, the reliable estimation on a given training dataset of predictive accuracy and stability of deep learning models (or of deep features used by external models) is still a gray area. The underlying risk is that of overfitting the training data, or worse to overfit the validation data if the labels are visible, which is typical when datasets are fully released at the end of a MI data science challenge. As the number of deep learning based studies in digital pathology is exponentially growing, we suggest that the progress of this field needs environments (*e.g.* DAPs) to prevent such pitfalls, especially if features distilled by the network are used as radiomics biomarkers to inform medical decision. Further, given an appropriate DAP, alternative model choices should be benchmarked on publicly available datasets, as usual in the general computer vision domain (*e.g.* ImageNet [26] or COCO [27]).

This study provides three main practical contributions to controlling for algorithmic bias and improving reproducibility of machine learning algorithms for digital pathology:

1. A Data Analysis Plan (DAP) specialized for digital pathology, tuned on the predictive evaluation of deep features, extracted by a network and used by alternative classification heads;
2. A benchmark dataset (HINT) of 53 727 tiles of histological images from 30 tissue types, derived from GTEx [28] for the recognition of tissue of origin of up to 30 classes;
3. An end-to-end machine learning framework (DAPPER) as a baseline environment for predictive models in digital pathology.

We first apply DAPPER to a set of classification experiments on 787 Whole Slide Images (WSIs) from GTEx. The framework (see Fig 1) is composed by (A) a preprocessing section to derive datasets from WSIs; (B) a 3-step machine learning pipeline with a data augmentation preprocessor, a backbone deep learning model, and an adapter extracting the deep features; (C) a downstream machine learning head, *i.e.* the task specific predictor. In our experiments, we analyze accuracy and feature stability in a multi-class setting of three different deep learning architectures (VGG, ResNet and Inception) as feature extractors and of three classifiers (a fully connected multilayer network, SVMs and Random Forests) as downstream machine learning head. This section is endowed with the DAP, i.e. a 10 × 5 CV (5-fold Cross Validation iterated 10 times). The 50 internal test sets are used to estimate a vector of metrics (with confidence intervals) that are used for model selection. In the fourth section (D) we finally provide an unsupervised data analysis based on the UMAP projection method and methods for feature exploration. The DAPPER software is available together with the HINT benchmark dataset.

**Fig 1.**
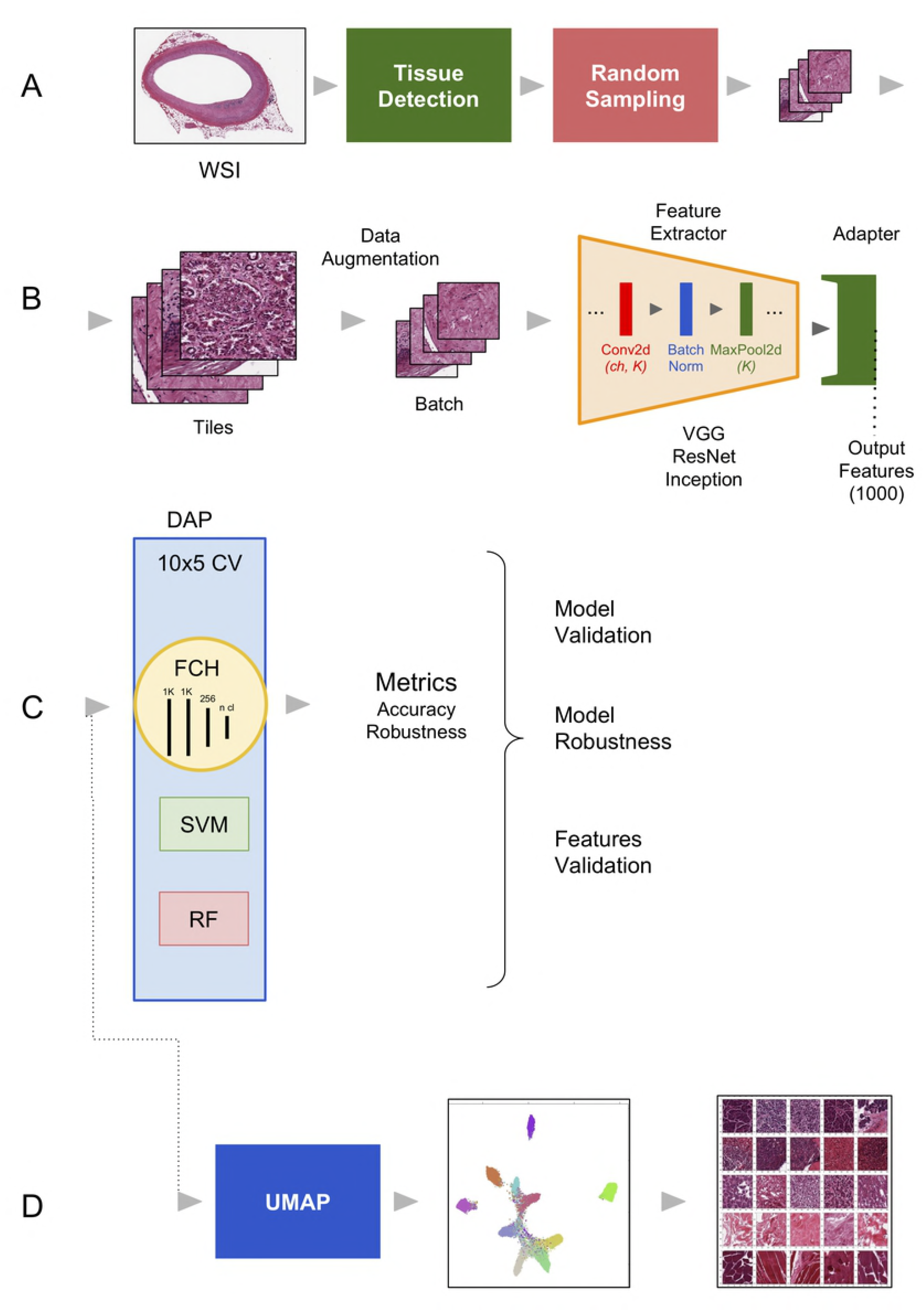
The DAPPER environment. Components: A) The WSI preprocessing pipeline; B) the deep learning backbone, to extract deep features; C) the Data Analysis Plan (DAP) for the machine learning models; and D) the UMAP module and other modules for unsupervised analysis.

Notably, the DAP estimates are provided in this paper only for the downstream machine learning head in section (C); whenever computational resources are available, the DAP can be expanded also to section (B). Here we kept as a separate problem the model selection exercise on the upstream deep learning backbone in order to clarify the change of perspective with respect to optimization of machine learning models in the usual training-validation setting.

As a second experiment, in order to study the DAPPER framework in a transfer learning condition, we use the deep features from the VGG model trained on HINT on the 1300 annotated tiles of the KIMIA24 dataset [29] to identify in this case the slide of origin (24 classes).

Previous work on classifying WSIs by means of neural networks was introduced by [29, 30], also with the purpose of distributing the two original datasets Kimia Path960 and Kimia Path24. Kimia Path24 consists of 24 WSIs chosen on purely visual distinctions. Babaie et al. [29] manually selected a total of 1325 binary patches with 40% overlap. On this dataset, amongst other two models based on Local Binary Patterns (LBP) and Bag-of-Words (BoVW), they applied two shallow convolutional neural networks (CNN), achieving at most 41.8% accuracy. Kimia Path960 contains 960 histopathological images belonging to 20 different WSIs that, again on visual clues, were used to represent different texture/pattern/staining types. Kumar et al. [30] replicated the same experiments as on Kimia Path24, *i.e.* LBP, BoVW and CNN. They applied AlexNet or VGG16, both pretrained on ImageNet, to extract deep features; instead of a classifier, accuracy was established by computing similarity distances between the 4096 features extracted. Also, Kiefer et al [31] explored the use of deep features from several pre-trained structures on Kimia24, controlling for the impact of transfer learning and finding an advantage of pre-trained networks against training from scratch. Alhindi and colleagues [32] instead analyze Kimia960 for slide of origin (20 slides preselected by visual ispection), and similarly to our study compare alternative classifiers as well as feature extraction models in a 3-fold CV setup. Our framework differs for the DAP structure and is originally applied to the larger HINT set without any visual pre-selection.

## Materials and methods

### Dataset

The images used to train the models were derived from the Genotype-Tissue Expression (GTEx) Study [28]. The study collects gene expression profiles and whole-slide images (WSIs) of 42 human tissues histologies used to investigate the relationship between genetic variation and tissue-specific gene expression in healthy individuals. To ensure that the collected tissues meet prescribed standard criteria, a Pathology Resource Center (PCR) validated each sample origin, content, integrity and target tissue (https://biospecimens.cancer.gov/resources/sops/). After sectioning and staining (H&E), tissue samples were scanned using a digital whole slide imaging system (Aperio) and stored in *.svs* format [33].

A custom Python script was used to download 787 WSIs through the Biospecimen Research Database (total size: 192 GB, average 22 WSIs for each tissue). The list of the downloaded WSIs is available in S1 Supplement Table S1.

A data preprocessing pipeline was developed to prepare the WSIs as training data (see Figure 2). The WSIs have a resolution of 0.275 *μm/pixel* (Magnification 40X) and variable dimension but the region interested by the tissue is only a portion of the WSI, which varies across the samples. Hence first we identified the region of the tissue in the image (see Fig 2), then we extracted at most 100 tiles (512 × 512 pixel) from the WSIs, by randomly sampling the tissue region. We applied the algorithm for the detection of the tissue region (see Fig 2) on each tile and rejected those where the portion of the tissue was below 85%. A total number of 53727 tiles was extracted, with a number of tiles per tissue varying between 59 (for Adipose - Visceral (Omentum)) and 2689 (for Heart - Left Ventricle).

**Fig 2.**
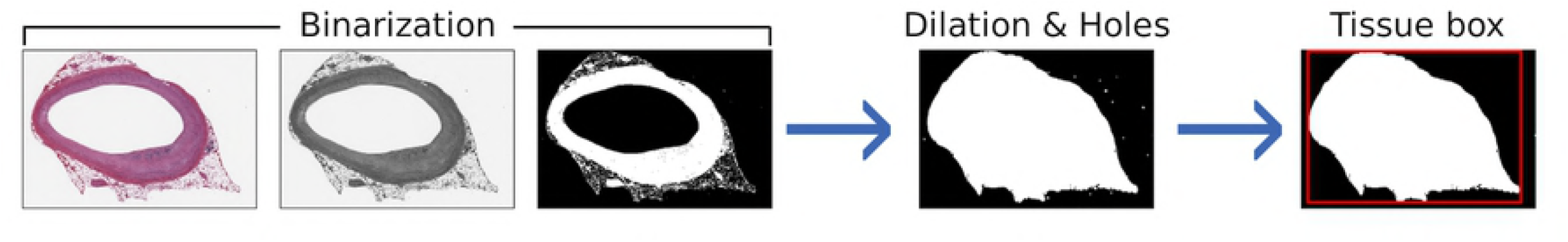
The tissue detection pipeline. The identification of the tissue bounding box is performed on the WSI thumbnail in three steps: a) Binarization of the grayscale image by applying Otsu’s thresholding; b) Binary dilation and filling of the holes; c) Selection of the biggest connected region as tissue region and computation of the vertex of the containing rectangle.

Four datasets (HINT5, HINT10, HINT20, HINT30) have been derived with increasing number of tissues for a total of 52 991 tiles (see S1 Supplement Table S2 for details). We refer to the four sets as the HINT collection, or the HINT dataset in brief. We choose the five tissues composing HINT5 starting from exploratory experiments, while the other datasets were composed including the tissues with higher number of tiles.

The class imbalance is accounted for by weighting the error on predictions. In detail, the weight *w* of the class *i* used in the cross entropy function is computed as: *w_i_* = *n_max_/n_i_*, where *n_max_* is the number of tiles in the class with more tiles and *n_i_* is the number of tiles in the class *i*.

Since image orientation should not be relevant for the tissue recognition, the tiles are randomly flipped (horizontally and vertically) and scaled, following a common practice in deep learning known as data augmentation [34]. Augmentation increases the generalization capabilities of the network preventing overfitting and is performed each time the tile is loaded, so that the resulting input image is different at each epoch. Such randomized transformations were found to increase concordance of prognostic accuracy of the deep learning SCNN architecture with that of baseline models based on combined molecular subtype and histologic grade [7]. In addition, the tile is cropped to a fixed size, which is dependent on the type of backbone network.

### Deep learning architectures and training strategies

We tested three backbone architectures commonly used in computer vision tasks (see Table 2):

1. VGG, version 19 with Batch Normalization (BN) layers [35];
2. ResNet, version 152 [36];
3. Inception, version 3 [37].

**Table 1.**
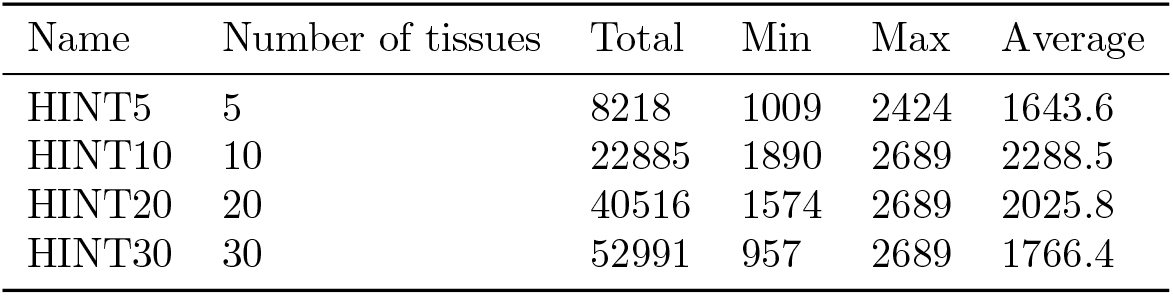
Summary of the HINT datasets. Total: total number of tiles composing the dataset; Min: number of tiles in the class with less samples; Max: number of tiles in the class with more samples; Average; average number of tiles for each class.

| Name | Number of tissues | Total | Min | Max | Average |
| --- | --- | --- | --- | --- | --- |
| HINT5 | 5 | 8218 | 1009 | 2424 | 1643.6 |
| HINT10 | 10 | 22885 | 1890 | 2689 | 2288.5 |
| HINT20 | 20 | 40516 | 1574 | 2689 | 2025.8 |
| HINT30 | 30 | 52991 | 957 | 2689 | 1766.4 |

**Table 2.** Backbone networks’ characteristics.

| Name | Output features | Parameters | Layers |
| --- | --- | --- | --- |
| VGG | 25088 | 20 035 392 | 64 |
| ResNet | 2048 | 58 143 808 | 465 |
| Inception | 2048 | 21 785 568 | 282 |

The feature extraction layers of each backbone network is obtained as the output of an end-to-end pipeline composed of four main blocks (see Fig 1, B):

1. Data augmentation: the input tiles are processed and assembled into batches;
2. Feature extraction: series of convolutional layers (Conv2d: with different number of channels and kernel size), normalization layers (Batch Norm) and pooling layers (MaxPool2d: with different kernel size) designed to fit with the backbone architecture. The number of output features of the Feature Extractor also depends on the backbone architecture;
3. Adapter: linear layer with 1000 output features, and the features of the backbone network as input;
4. Machine Learning Head: task specific model taking the 1000 Adapter features as input and providing predicted tissue labels (or a clinical endpoint) as output.

As predictive models, we used a linear Support Vector Machine (SVM) with regularization parameter C set to 1, a Random Forest (RF) classifier with 500 trees (both implemented in *scikit-learn,* v0.19.1) and a fully connected head (FCH), *i.e.* a series of fully connected layers. Inspired by [7], our FCH consists of four dense layers with 1000, 1000, 256 and # tissue classes nodes, respectively. The feature extraction block was initialized with the weights already trained on the ImageNet dataset [26], provided by *PyTorch* (v0.4.0) and freezed. Training also the weights of the feature extraction block improves accuracy (see S1 Supplement Table S3). However, these results were not validated rigorously within the DAP and therefore they not are not claimed as generalized in this study.

For the optimization of the other weights (Adapter and FCH) we used the Adam algorithm [38] with the learning rate set to 10^−5^ and fixed for the whole training. We used the cross entropy as loss function, which is appropriate for multi-class models.

The strategy to optimize the learning rate was selected based on results of a preparatory study with the VGG network and HINT5. The strategy approach with fixed learning rate achieved the best results (see S1 Supplement Table S4) and was therefore adopted in the rest of the study.

### Data Analysis Plan

Following the rigorous model validation techniques proposed by the MAQC projects [23, 24], we adopted a DAP to assess the validity of the features extracted by the networks, namely a 10×5-fold cross-validation (CV) schema. The input dataset is first partitioned with 80% of samples used as training set for model development and 20% as held-out test set for model evaluation. The training set further undergoes a 5-fold CV iterated 10 times resulting in a ranked feature list and a set of metrics, including accuracy (ACC) and the Matthews Correlation Coefficient (MCC) in its multiclass generalization [39, 40]. At each CV iteration, features are ranked by ANOVA F-score and the classifier is trained on an increasing number of ranked features (namely: 10%, 25%, 50%, 100% of the total number of features). A list of top-ranked features is obtained by Borda aggregation of the ranked lists, for which we also compute the Canberra stability with a computational framework designed for sets of ranked biomarker lists [25]. The overall performance of the predictive model is evaluated across all iterations in terms of average MCC and ACC with 95% Studentized bootstrap confidence intervals (CI), and on the test set in terms of MCC.

As a sanity check to avoid unwanted selection bias effects, the DAP is repeated stochastically scrambling the training set labels (*random labels* mode: in absence of selection bias, the no-information error rate is expected) or by randomly ranking features before building models (*random ranking* mode: in presence of pools of highly correlated variables, top features can be interchanged with others, possibly of higher biological interest).

### Experiments on HINT

We designed a set of experiments to provide indications about the optimal backbone architecture, while keeping fixed the other hyper-parameters. In particular we set batch size (32) and number of epochs (50), high enough to let the network converge: we explored increasing numbers of epochs (10, 30, 50, 100) and as we observed that the loss stabilizes after about 35 epochs, we set the number of epochs to 50. First, we compared the three backbone architectures on the smallest dataset HINT5, with fixed learning rate. Both VGG and ResNet architectures achieved good results, far outperforming Inception. In further analyses we thus restricted to use VGG and ResNet as feature extractors and validated performance and features with the DAP. The same process was adopted on HINT10 and HINT20. An experiment with 30 tissues has also been performed. Results are listed in S1 Supplement Table S5. A summary of the experiments on backbone models is reported in Table 3.

**Table 3.** Summary of experiments with the backbone models.

| Experiment | # tissues | Feature extractor | VGG Version |
| --- | --- | --- | --- |
| VGG-5 | 5 | VGG | 19+BN |
| ResNet-5 | 5 | ResNet | 152 |
| Inception-5 | 5 | Inception | 3 |
| VGG-10 | 10 | VGG | 19+BN |
| ResNet-10 | 10 | ResNet | 152 |
| VGG-20 | 20 | VGG | 19+BN |
| ResNet-20 | 20 | ResNet | 152 |

### UMAP Analysis

The UMAP multidimensional analysis method [41] was applied to project the features extracted onto a bi-dimensional space, using the Python *umap* module.

### Implementation

All the code used for image processing and network training is written in Python (v3.6) and is available as a collection of Jupyter notebooks at https://gitlab.fbk.eu/histology_DL/Tissue_classification.

In addition to the general scientific libraries for Python, the scripts for the creation and training of the networks are based on `PyTorch`; the backbone networks are implemented in *torchvision.* The library for processing histological images (available at https://gitlab.fbk.eu/histology_DL/histo_lib) is based on *OpenSlide* and *scikit-image.*

The computations were performed on Microsoft Azure Virtual Machines with 4 NVIDIA K80 GPUs, 24 Intel Xeon E5-2690 cores and 256 GB RAM.

## Results

Results of two digital pathology tasks in the DAPPER framework are listed in Table 4 for Matthews Correlation Coefficient (MCC) and Table 5 for Accuracy (ACC), respectively. See also Fig 3 (a-b-c) for a comparison of MCC in internal cross-validation with external validation.

**Table 4.** DAPPER results for each deep learning backbone and classifier head pair on HINT datasets and KIMIA24: average cross-validation MCC with 95% Confidence Intervals (CI) and MCC on test set.

| Experiment | FCH |  | SVM |  | RF |  |
| --- | --- | --- | --- | --- | --- | --- |
|  | MCC (CI) | MCC Test | MCC (CI) | MCC Test | MCC (CI) | MCC Test |
| VGG-5 | 0.841 (0.838, 0.843) | 0.820 | 0.786 (0.783, 0.789) | 0.777 | 0.750 (0.748, 0.753) | 0.747 |
| ResNet-5 | **0.879 (0.877, 0.881)** | **0.883** | 0.852 (0.850, 0.854) | 0.840 | 0.829 (0.827, 0.832) | 0.849 |
| VGG-10 | **0.896 (0.894, 0.897)** | **0.894** | 0.861 (0.859, 0.862) | 0.866 | 0.889 (0.888, 0.891) | 0.886 |
| ResNet-10 | 0.857 (0.856, 0.859) | 0.860 | 0.825 (0.824, 0.827) | 0.832 | 0.845 (0.843, 0.847) | 0.850 |
| VGG-20 | 0.771 (0.770, 0.772) | 0.774 | 0.729 (0.727, 0.730) | 0.731 | 0.761 (0.760, 0.762) | 0.766 |
| ResNet-20 | 0.756 (0.754, 0.757) | 0.757 | **0.788 (0.787, 0.789)** | **0.792** | 0.738 (0.737, 0.739) | 0.738 |
| VGG-KIMIA24 | 0.317 (0.306, 0.327) | 0.207 | 0.446 (0.439, 0.454) | 0.409 | **0.457 (0.449, 0.465)** | **0.409** |

**Table 5.**
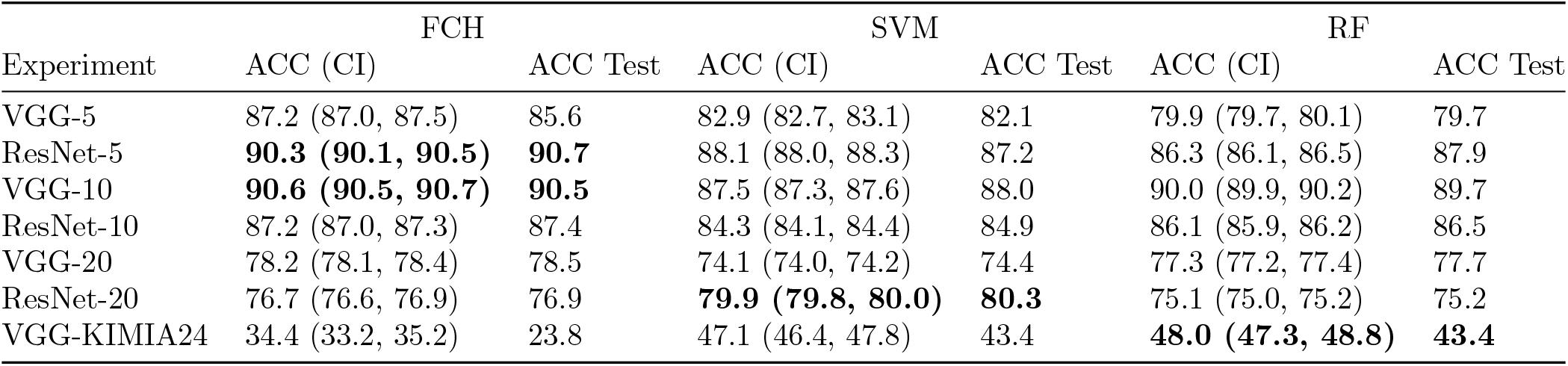
DAPPER results for each deep learning backbone and classifier head pair on HINT datasets and KIMIA24: average cross-validation accuracy with 95% Confidence Intervals (CI) and accuracy on test set.

**Fig 3.**
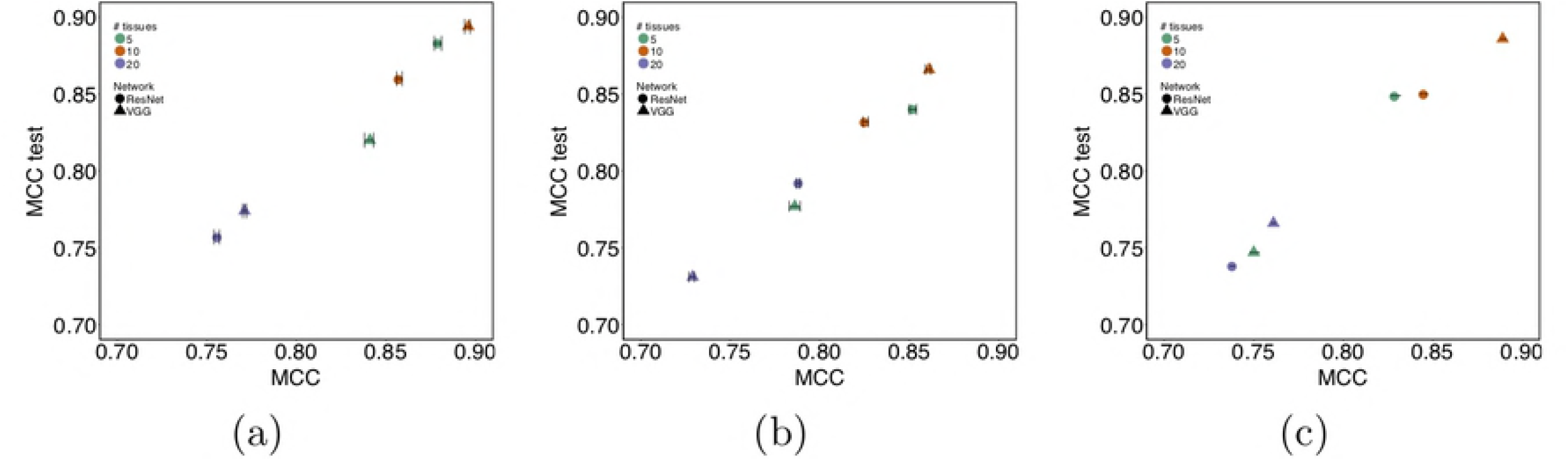
Comparison of DAPPER cross-validation (MCC) vs external validation (MCC Test) performance for each classifier. (a) FCH; (b) SVM; (c) RF.

All backbone-head pairs on HINT have MCC>0.7 with narrow CIs, with estimates from internal validation close to performance on held-out test (Fig 3). Agreement of internal estimates with values on test set is a good indicator of generalization and potential for reproducibility. All models reached their top MCC accuracy with 1000 features. On HINT5 and HINT10, the FCH neural network performs better than SVMs and RF. As expected, MCC ranged close to 0 for random labels; random ranking for increasing feature set sizes reached top MCC only for all features (tested for SVMs, results not shown).

The most accurate models both for training estimates and in validation were the (ResNet,FCH) pair with MCC=0.883 on HINT5, the (VGG,FCH) pair on HINT10, the (Resnet,SVM) on HINT20. Results on HINT30 are detailed in S1 Supplement Table S5; on test, VGG-HINT30 reaches accuracy ACC= 61.8% and MCC=0.61. Performance decreases for more complex multiclass problems. Notably the difficulty of the task is also complicated by tissue classes that are likely to have similar histological patterns, such as misclassification of Esophagus-Muscularis (ACC: 72.1%) with Esophagus-Mucosa (ACC: 53.2%), or the two Heart tissue subtypes or the 58 Ovary(ACC: 68.3%) tiles predicted as Uterus (ACC: 72.8%). The complete confusion matrix for Resnet with SVMs on HINT20 is reported in Fig 4.

**Fig 4.**
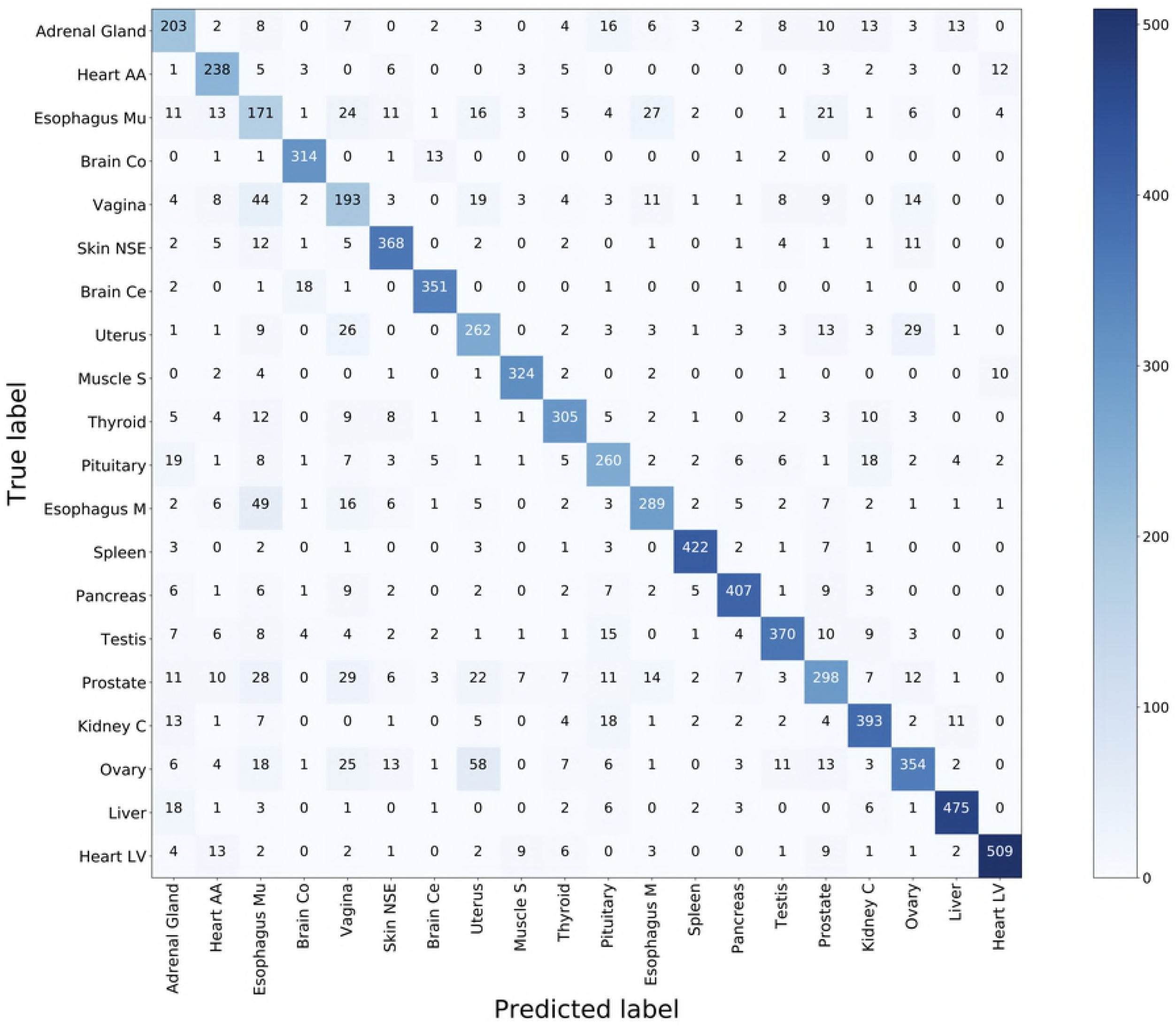
Confusion matrix for SVM on ResNet for HINT20. Entries are counts of predicted vs true label for the 20 tissues. Heart AA: Heart-Atrial Appendage; Heart LV: Heart-Left Ventricle; Esophagus Mu: Esophagus-Mucosa; Esophagus M: Esophagus-Muscularis; Brain Co: Brain-Cortex; Brain Ce: Brain-Cerebellum; Skin NSE: Skin-Not Sun Exposed (Suprapubic); Muscle S: Muscle-Skeletal; Kidney C: Kidney-Cortex.

In the second experiment, we used the VGG with deep features from the VGG from GTEx on the Kimia Path24 (KIMIA24) dataset [29], where the task is identifying slide of origin (24 classes). In the DAPPER framework, classifiers were trained on 1060 annotated tiles and validated on 265 unseen ones. Regardless of difference in image types, VGG-KIMIA24 with both RF and SVM heads with ACC=43.4% (see Table 5), improving on published results (ACC=41.8%; [29]). The FCH had lower accuracy, but it should be taken into account that deep features and classifiers were trained on 1060 images only, compared to the 40513 used by Babaie and colleagues.

It is worth noting that transfer learning from ImageNet to HINT restricts training to the Adapter and Fully Connected Head blocks. In one-shot experiments, MCC further improves when the whole feature extraction block is retrained (see S1 Supplement Table S3). However, the result still needs to be consolidated by extending the DAP also to the training or retraining of the deep learning backbone to check for actual generalization. The Canberra stability indicator was also computed for all the experiments, with minimal median stability for ResNet-HINT20 (Fig 5).

**Fig 5.**
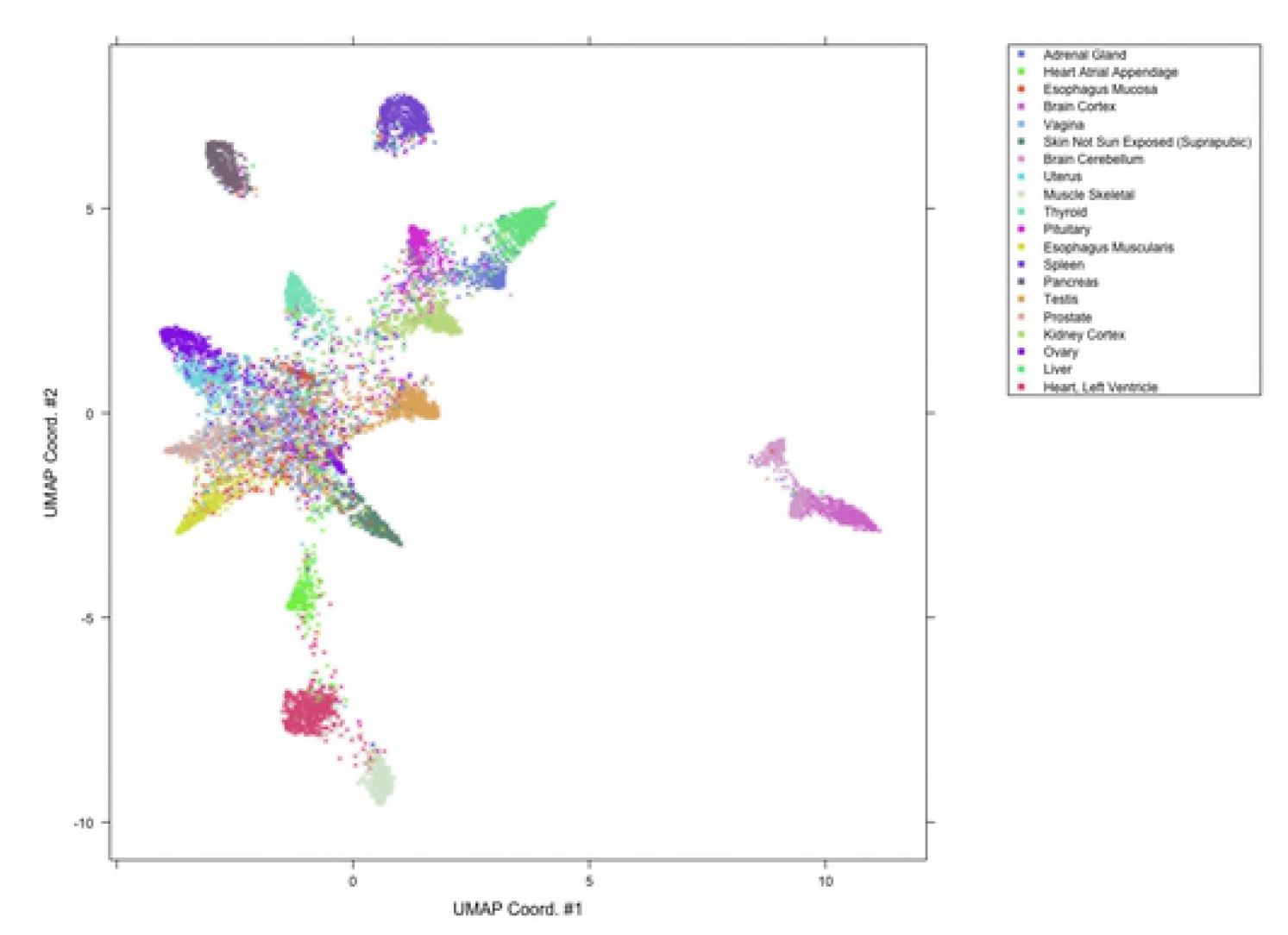
Canberra stability indicator. For each architecture, a set of deep feature lists is generated, one list for each internal run of training in the nested cross-validation schema, each ranked with KBest. Canberra stability is computed as in [25]: lower stability is better.

## The HINT Benchmark Dataset

As a second contribution of this study, we are making available the HINT dataset, generated by the first section of tools in the DAPPER framework, as a benchmark dataset for validating machine learning models in digital pathology. The HINT dataset is currently composed of 53727 tiles at 512 × 512 resolution, based on histology from GTEx. HINT can be easily expanded to over 78000 tiles, as for this study we used a fraction of the GTEx images and at most 100 tiles from each WSI were extracted. Digital pathology still misses a universally adopted dataset to compare deep learning models as already established in vision (e.g. ImageNet for image classification, COCO for image and instance segmentation). Several initiatives for a “BioImageNet” will eventually improve this scenario. Histology data are available in the generalist repository Image Data Resource (IDR) [42, 43]. Further, the International Immuno-Oncology Biomarker Working Group in Breast Cancer and the MAQC Society have launched a collaborative project to develop data resources and quality control schemes on machine Learning algorithms to assess TILs in Breast Cancer.

HINT is conceptually similar to KIMIA24. However, HINT inherits from GTEx more variability in terms of sample characteristics, validation of donors and additional access to molecular data. Further, we used a random sampling approach to process tiles excluding background and minimize human intervention in the choice and preparation of the images.

### Deep features

We applied an unsupervised projection on all the features extracted by VGG and ResNet networks on all tissues tasks. In the following, we discuss an example for features extracted by VGG on the HINT20 task, displayed as UMAP projection in Fig 6; points are coloured for 20 tissue labels. The UMAP displays for the other tasks are available in S1 Supplement Fig S1-S4.

**Fig 6.**
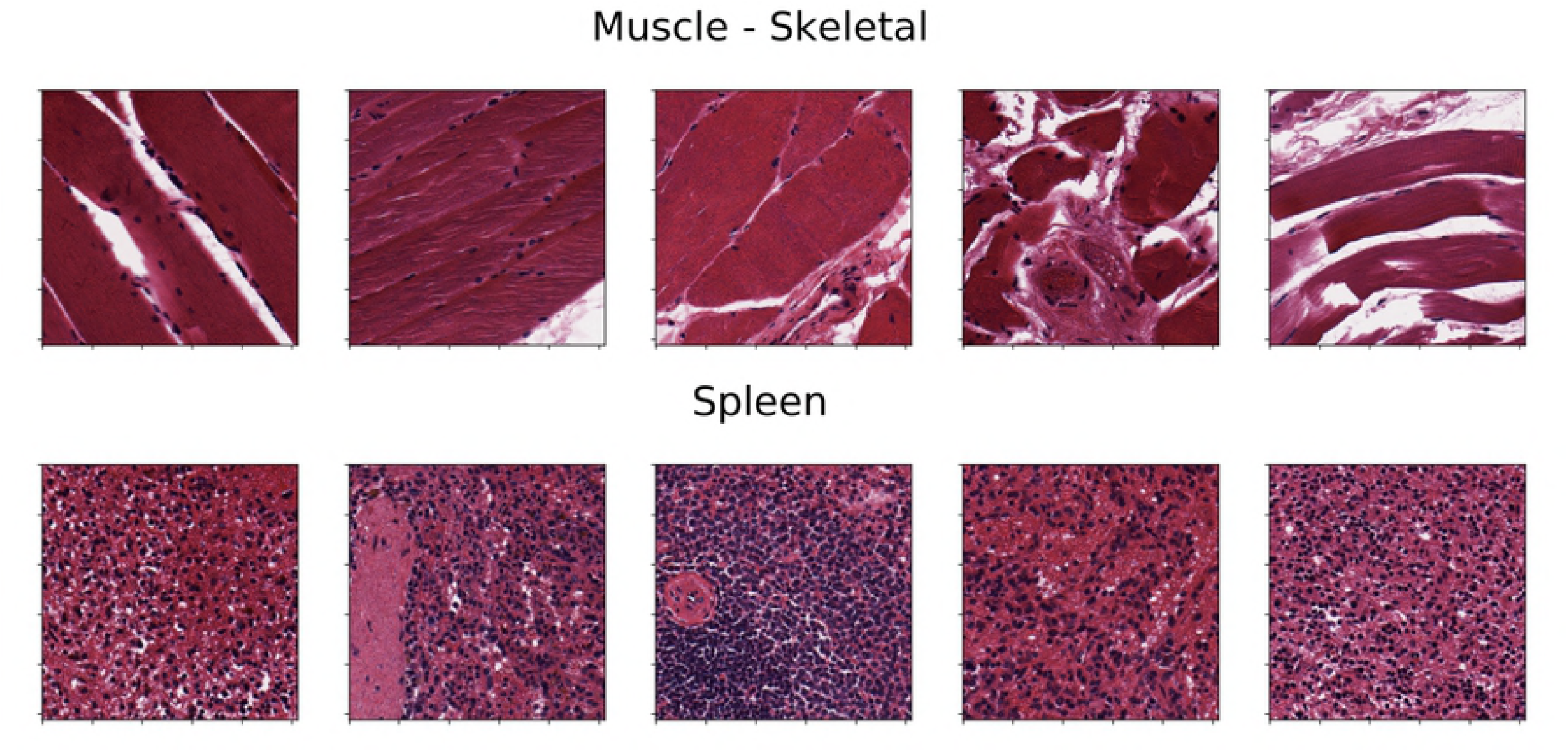
UMAP projection of validation set for VGG-20 task.

The UMAP display is in agreement with the count distributions in the confusion matrix (Fig 4). The deep learning embedding separates well a set of histology types, including: Muscle - Skeletal (ACC: 93.4%), Spleen (ACC: 94.6%), Pancreas (ACC: 87.9%), Brain - Cortex (ACC: 94.3%) and Brain - Cerebellum (ACC: 93.4%), Heart - Left Ventricle (ACC: 90.1%) and Heart - Atrial Appendage (ACC: 84.7%) group into distinct clusters.

The distributions of the activations for the top-3 deep features of the VGG-HINT10 backbone are displayed in S1 Supplement Fig S5. The top ranked deep feature (#668) is clearly selective for Spleen.

The projection also shows an overlapping for tissues such as Ovary and Uterus, or Vagina and Esophagus - Mucosa, or the two Esophagus histotypes, consistently with the confusion matrix (Fig 4).

Examples of five tiles from two well separated clusters, Muscle-Skeletal (ACC: 93.4%) and Spleen (ACC: 94.6%), are displayed in Fig 7. Tiles from five clusters partially overlapping in the neural embedding and mislabeled in both the VGG-20 and ResNet-20 embeddings with SVMs (Uterus ACC=72.8%, Ovary ACC=68.3%, Esophagus-Mucosa ACC=53.2%, Esophagus-Muscularis ACC=72.1%, Vagina ACC=59.0%) are similarly visualized in Fig 8.

**Fig 7.**
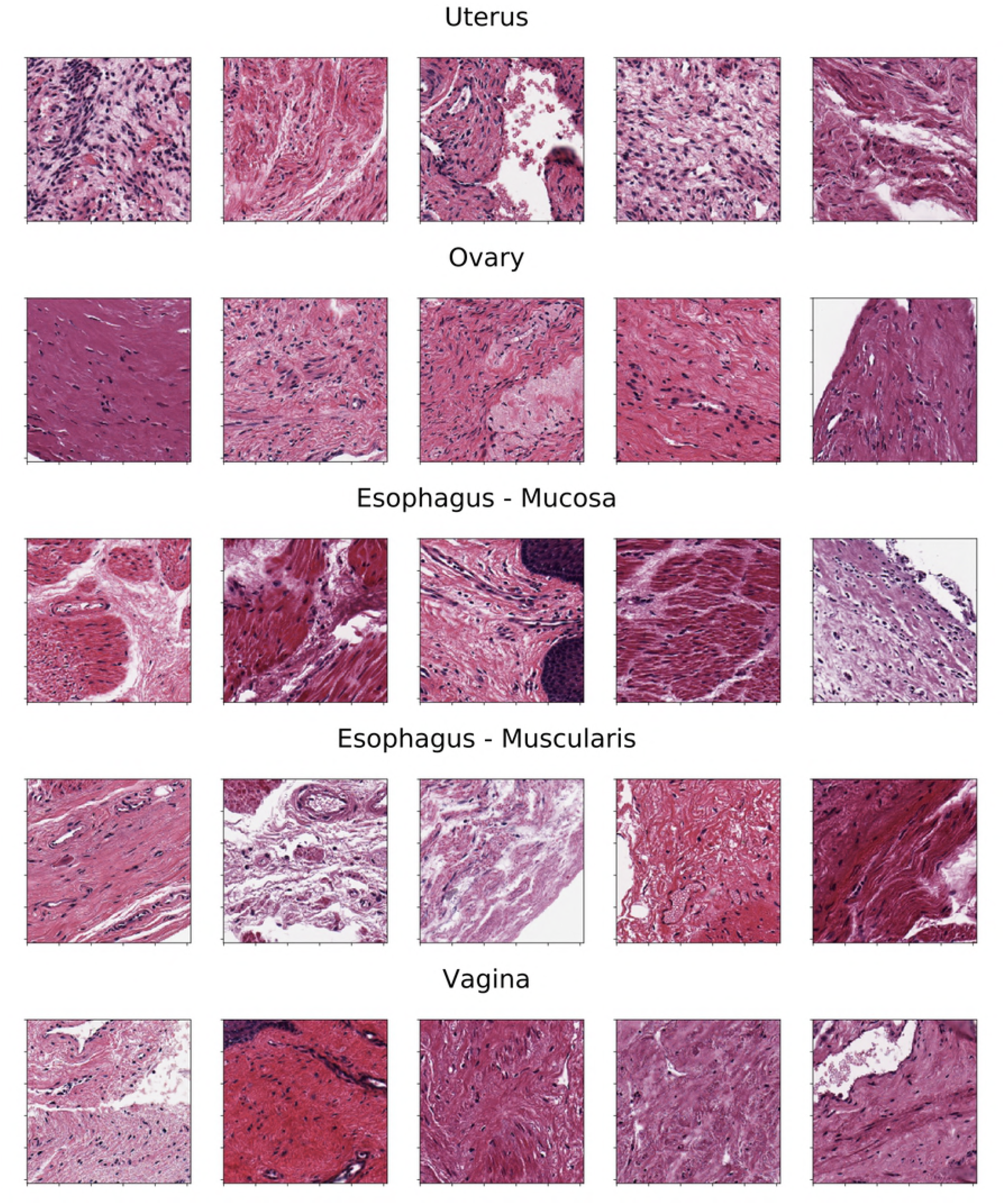
Representative tiles from two well-separated clusters observed on the UMAP embedding of VGG-20.

**Fig 8.**
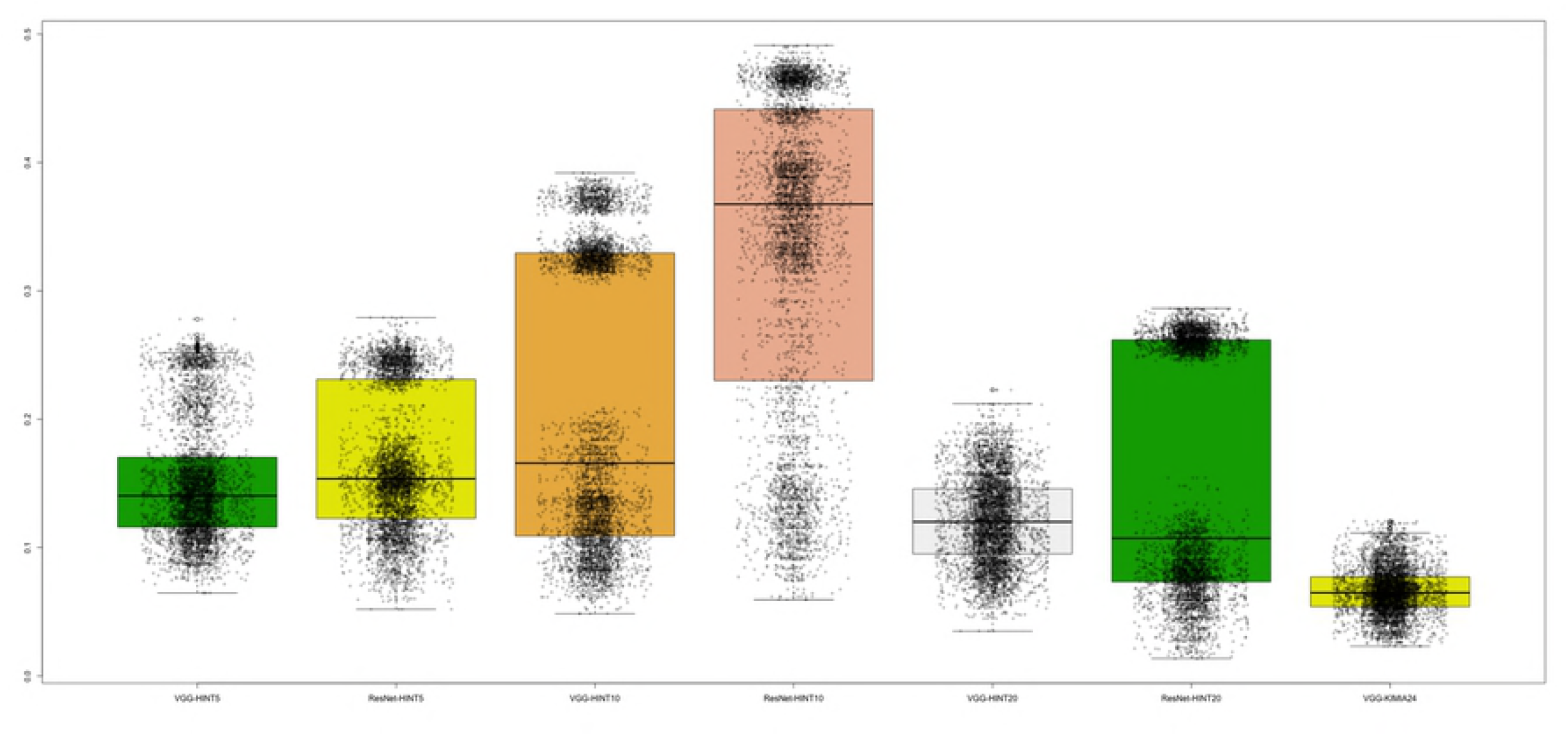
Samples of mislabeled tiles from five tissues partially overlapping in the VGG-20 UMAP embedding.

While the aim of this paper is to introduce a framework for honest comparison of models that will be used for clinical purposes rather than fine-tuning accuracy in this experiment, it is evident that these tiles have morphologies that are hard to classify. This challenge requires more complex models (e.g. ensembles) and a structured output labeling, already applied in dermatology [2].

Further, we are exploring the combination of DAPPER with image analysis packages, such as HistomicsTK (https://digitalslidearchive.github.io/HistomicsTK/ or CellProfiler [44], to extract features useful for interpretation and feedback from pathologists.

## Conclusions

Digital pathology would greatly benefit from the adoption of machine learning, shifting human assessment of histology to higher quality, non-repetitive tasks. Unfortunately, there is no fast, easy route to improve reproducibility of automated analysis. The adoption of the DAP clearly sets in a computational aggravation not usually considered for image processing exercises. However, this is an established practice with massive omics data [21], and reproducibility by design can handle secondary results useful for diagnostics and for interpretation.

We designed the DAPPER framework as a tool for evaluating accuracy and stability of deep learning models, currently only backbone elements in a sequence of processing steps, and possibly in the future end-to-end solutions. We choose as test domain H&E stained WSIs for prediction of tissue of origin, which is not a primary task for trained pathologists, but a reasonable benchmark for machine learning methods. Also, we are aware that tissue classification is only a step in real digital pathology applications. Mobadersany and colleagues [7] used a deep learning classifier to score and visualize risk on the WSIs. Similarly, deep learning tile classification may be applied to quantify histological differences in association to a genomic pattern, e.g. a specific mutation or a high-dimensional protein expression signature. In this vision, the attention to model selection supported by our framework is a prerequisite for developing novel AI algorithms for digital pathology, e.g. for analytics over TILs.

Although we are building on deep learning architectures known for applications on generic images, they adapted well to WSIs in combination with established machine learning models (SVMs, RF); we expect that large scale bioimaging resources will give the chance of improving the characterization of deep features, as already emerged with the HINT dataset that we are providing as public resource. In this direction, we plan to release the network weights of the backbone DAPPER models that are optimized for histopathology as alternative pretrained weights for digital pathology, similarly to those for the ImageNet dataset and available in torchvision.

## Supporting information

**S1 Supplement. Supporting tables and figures.**

## Acknowledgments

Cloud computing was funded by the Azure Research grant “Deep Learning for Precision Medicine”, endowed to CF. We derived the HINT test dataset from the Genotype-Tissue Expression (GTEx) Project, supported by the Common Fund of the Office of the Director of the National Institutes of Health, and by NCI, NHGRI, NHLBI, NIDA, NIMH, and NINDS (data downloaded from the GTEx Portal on *05/10/18*). The authors thank H. Tizhoosh too for availability of Kimia Path24 dataset.

## References

1. Lu L, Zheng Y, Carneiro G, Yang L. Deep Learning and Convolutional Neural Networks for Medical Image Computing. Springer; 2017.

2. Esteva A, Kuprel B, Novoa RA, Ko J, Swetter SM, Blau HM, et al. Dermatologist-level classification of skin cancer with deep neural networks. Nature. 2017;542(7639):115.

3. Litjens G, Kooi T, Bejnordi BE, Setio AAA, Ciompi F, Ghafoorian M, et al. A survey on deep learning in medical image analysis. Medical Image Analysis. 2017;42:60–88.

4. Komeda Y, Handa H, Watanabe T, Nomura T, Kitahashi M, Sakurai T, et al. Computer-aided diagnosis based on convolutional neural network system for colorectal polyp classification: preliminary experience. Oncology. 2017;93(Suppl. 1):30–34.

5. Korbar B, Olofson AM, Miraflor AP, Nicka CM, Suriawinata MA, Torresani L, et al. Deep learning for classification of colorectal polyps on whole-slide images. Journal of Pathology Informatics. 2017;8.

6. Ciompi F, Chung K, Van Riel SJ, Setio AAA, Gerke PK, Jacobs C, et al. Towards automatic pulmonary nodule management in lung cancer screening with deep learning. Scientific Reports. 2017;7:46479.

7. Mobadersany P, Yousefi S, Amgad M, Gutman DA, Barnholtz-Sloan JS, Velázquez Vega JE, et al. Predicting cancer outcomes from histology and genomics using convolutional networks. Proceedings of the National Academy of Sciences. 2018;115(13):E2970–E2979.

8. Bychkov D, Linder N, Turkki R, Nordling S, Kovanen PE, Verrill C, et al. Deep learning based tissue analysis predicts outcome in colorectal cancer. Scientific Reports. 2018;8(1).

9. Sharma H, Zerbe N, Klempert I, Hellwich O, Hufnagl P. Deep convolutional neural networks for automatic classification of gastric carcinoma using whole slide images in digital histopathology. Computerized Medical Imaging and Graphics. 2017;61:2–13.

10. Paeng K, Hwang S, Park S, Kim M. A unified framework for tumor proliferation score prediction in breast histopathology. Deep Learning in Medical Image Analysis and Multimodal Learning for Clinical Decision Support. 2017; p. 231–239.

11. Basavanhally ea A. Multi-field-of-view framework for distinguishing tumor grade in ER+ breast cancer from entire histopathology slides. Biomed Eng. 2013;60:2089–2099.

12. Denkert C, Wienert S, Poterie A, Loibl S, Budczies J, Badve S, et al. Standardized evaluation of tumor-infiltrating lymphocytes in breast cancer: results of the ring studies of the international immuno-oncology biomarker working group. Modern Pathology. 2016;29(10):1155.

13. Mina M, Boldrini R, Citti A, Romania P, D’Alicandro V, De Ioris M, et al. Tumor-infiltrating T lymphocytes improve clinical outcome of therapy-resistant neuroblastoma. Oncoimmunology. 2015;4(9):e1019981.

14. Salgado R, Sherene L. Tumour infiltrating lymphocytes in breast cancer: increasing clinical relevance. The Lancet Oncology. 2018;19(1):3–5.

15. Stovgaard ES, Nielsen D, Hogdall E, Balslev E. Triple negative breast cancer–prognostic role of immune-related factors: a systematic review. Acta Oncologica. 2018;57(1):74–82.

16. Shibutani M, Maeda K, Nagahara H, Fukuoka T, Iseki Y, Matsutani S, et al. Tumor-infiltrating Lymphocytes Predict the Chemotherapeutic Outcomes in Patients with Stage IV Colorectal Cancer. In Vivo. 2018;32(1):151–158.

17. Tajbakhsh N, Shin JY, Gurudu SR, Hurst RT, Kendall CB, Gotway MB, et al. Convolutional neural networks for medical image analysis: Full training or fine tuning? IEEE transactions on medical imaging. 2016;35(5):1299–1312.

18. Kieffer B, Babaie M, Kalra S, Tizhoosh H. Convolutional Neural Networks for Histopathology Image Classification: Training vs. Using Pre-Trained Networks. arXiv preprint arXiv:171005726. 2017;.

19. Ioannidis JP, Allison DB, Ball CA, Coulibaly I, Cui X, Culhane AC, et al. Repeatability of published microarray gene expression analyses. Nature genetics. 2009;41(2):149.

20. Baker M. 1,500 scientists lift the lid on reproducibility. Nature News. 2016;533(7604):452.

21. Shi L, Kusko R, Wolfinger RD, Haibe-Kains B, Fischer M, Sansone SA, et al. The international MAQC Society launches to enhance reproducibility of high-throughput technologies. Nature Biotechnology. 2017;35(12):1127.

22. Wilkinson MD, Dumontier M, Aalbersberg IJ, Appleton G, Axton M, Baak A, et al. The FAIR Guiding Principles for scientific data management and stewardship. Scientific data. 2016;3.

23. The MicroArray Quality Control (MAQC) Consortium. The MAQC-II Project: A comprehensive study of common practices for the development and validation of microarray-based predictive models. Nature Biotechnology. 2010;28(8):827–838.

24. The SEQC/MAQC-III Consortium. A comprehensive assessment of RNA-seq accuracy, reproducibility and information content by the Sequence Quality Control consortium. Nature Biotechnology. 2014;32:903–914.

25. Jurman G, Merler S, Barla A, Paoli S, Galea A, Furlanello C. Algebraic stability indicators for ranked lists in molecular profiling. Bioinformatics. 2008;24(2):258–264.

26. Deng J, Dong W, Socher R, Li LJ, Li K, Fei-Fei L. Imagenet: A large-scale hierarchical image database. In: Computer Vision and Pattern Recognition, 2009. CVPR 2009. IEEE Conference on. IEEE; 2009. p. 248–255.

27. Lin TY, Maire M, Belongie S, Hays J, Perona P, Ramanan D, et al. Microsoft coco: Common objects in context. In: European conference on computer vision. Springer; 2014. p. 740–755.

28. The GTEx Consortium. The genotype-tissue expression (GTEx) project. Nature genetics. 2013;45(6):580.

29. Babaie M, Kalra S, Sriram A, Mitcheltree C, Zhu S, Khatami A, et al. Classification and Retrieval of Digital Pathology Scans: A New Dataset. In: CVMI Workshop@ CVPR; 2017.

30. Kumar MD, Babaie M, Zhu S, Kalra S, Tizhoosh H. A Comparative Study of CNN, BoVW and LBP for Classification of Histopathological Images. arXiv preprint arXiv:171001249. 2017;.

31. Kieffer B, Babaie M, Kalra S, Tizhoosh HR. Convolutional Neural Networks for Histopathology Image Classification: Training vs. Using Pre-Trained Networks. CoRR. 2017;abs/1710.05726.

32. Alhindi TJ, Kalra S, Ng KH, Afrin A, Tizhoosh HR. Comparing LBP, HOG and Deep Features for Classification of Histopathology Images. arXiv preprint arXiv:180505837. 2018.

33. Carithers LJ, Ardlie K, Barcus M, Branton PA, Britton A, Buia SA, et al. A novel approach to high-quality postmortem tissue procurement: the GTEx project. Biopreservation and biobanking. 2015;13(5):311–319.

34. Wang J, Luis P. The effectiveness of data augmentation in image classification using deep learning. Technical Report; 2017.

35. Simonyan K, Zisserman A. Very deep convolutional networks for large-scale image recognition. arXiv preprint arXiv:14091556. 2014.

36. He K, Zhang X, Ren S, Sun J. Deep residual learning for image recognition. In: Proceedings of the IEEE conference on computer vision and pattern recognition; 2016. p. 770–778.

37. Szegedy C, Vanhoucke V, Ioffe S, Shlens J, Wojna Z. Rethinking the inception architecture for computer vision. In: Proceedings of the IEEE Conference on Computer Vision and Pattern Recognition; 2016. p. 2818–2826.

38. Kinga D, Adam JB. A method for stochastic optimization. In: International Conference on Learning Representations (ICLR); 2015.

39. Matthews BW. Comparison of the predicted and observed secondary structure of T4 phage lysozyme. Biochimica et Biophysica Acta. 1975;405(2):442–451.

40. Baldi P, Brunak S, Chauvin Y, et al. Assessing the accuracy of prediction algorithms for classification: an overview. Bioinformatics. 2000;16(5):412–424.

41. McInnes L, Healy J. UMAP: Uniform Manifold Approximation and Projection for Dimension Reduction; 2018.

42. Williams E, Moore J, Li SW, Rustici G, Tarkowska A, Chessel A, et al. Image Data Resource: a bioimage data integration and publication platform. Nature methods. 2017;14(8):775.

43. Nirschl JJ, Janowczyk A, Peyster EG, Frank R, Margulies KB, Feldman MD, et al. A deep-learning classifier identifies patients with clinical heart failure using whole-slide images of H&E tissue. PloS one. 2018;13(4):e0192726.

44. Carpenter AE, Jones TR, Lamprecht MR, Clarke C, Kang IH, Friman O, et al. CellProfiler: image analysis software for identifying and quantifying cell phenotypes. Genome biology. 2006;7(10):R100.

